# PocketX: Preference Alignment for Protein Pockets Design through Group Relative Policy Optimization

**DOI:** 10.64898/2025.12.28.696754

**Authors:** Yuliang Fan, Zaikai He, Bin Li, Bin He, Mingshu Zhang, Jian Zhang, Haicang Zhang

## Abstract

Designing protein pockets that target specific ligands is crucial for drug discovery and enzyme engineering. Although deep generative models show promise in proposing high-quality pockets, they are usually trained purely to match the data distribution and therefore overlook key biophysical properties, such as binding affinity, expression, and solubility, that ultimately determine developability and success. We introduce PocketX, an online reinforcement learning framework that explicitly aligns a generative model with desired biophysical properties. The framework first trains a base model that co-designs pocket structures and sequences conditioned on a target ligand, and then fine-tunes this model with Group Relative Policy Optimization (GRPO) to reward the desired attributes. Because GRPO employs group-relative rewards, it produces lower-variance policy updates, resulting in more stable and efficient learning than competing alignment strategies. Evaluated on the CrossDocked2020 benchmark, PocketX surpasses existing methods in metrics such as binding energy and evolutionary plausibility. Ablation studies further show that GRPO outperforms alternative alignment strategies, including Direct Preference Optimization (DPO), confirming GRPO’s effectiveness for biophysical property alignment.

## 1 Introduction

Designing protein pockets that bind specific ligands is fundamental to advances in drug discovery [37], clinical diagnostics [26], enzyme engineering [5], and biosensor development [44]. Existing computational approaches fall into two main categories: physics-based and template-based. Physics-based algorithms, such as PocketOptimizer [20, 35], perform combinatorial searches over sequence space to minimize calculated binding free energy. Template-based methods [3, 45, 25, 18] construct pockets by assembling structural motifs around the target ligand while enforcing specific hydrogen-bonding patterns. Despite their successes in particular cases, both strategies are constrained by limited accuracy and substantial computational cost.

Recent breakthroughs in deep generative modeling for languages [39, 21] and images [33, 29] are now reshaping protein pocket design. For example, FAIR [46] and PocketGen [47] utilize graph-based neural networks to predict the pocket structures and sequences directly, whereas RFdiffusion [42] and its variant RFdiffusion All-Atom (RFdiffusionAA) [14] leverage diffusion-based generative models to sample the pocket sequence and structures. Because these methods are trained primarily to reproduce the statistical distribution of existing data, they often neglect critical biophysical properties—such as binding affinity, expression level, and solubility—that ultimately determine developability and success in practice [24, 7].

To move beyond simple distribution matching, reinforcement learning (RL) provides a principled way to steer generation toward task-specific objectives. In molecular design, Direct Preference Optimization (DPO)–based methods have already improved antibody and small-molecule design [49, 28, 10] with more desired properties such as binding energy and stability. More recently, DeepSeek’s Group Relative Policy Optimization (GRPO) [11, 32]—an online RL algorithm originally developed for autoregressive language models—has been shown to surpass DPO and PPO [31] in both training stability and sample efficiency [11]. GRPO [11, 32] has since been generalized beyond autoregressive language modeling to diffusion- and flow-based generative models such as DanceGRPO [43] and FlowGRPO [16] and also achieves similar performance gains.

Motivated by these advances, we propose PocketX, a deep generative model for protein pocket design that leverages GRPO to steer the designed pockets toward desired biophysical properties. PocketX first trains a hybrid continuous–discrete diffusion model for pocket structure and sequence co-generation, using AlphaFold3-style architectures as the score network [1]. The resulting base model is then fine-tuned with GRPO. Physics-based rewards, such as AutoDock Vina docking scores [38, 8], promote high binding affinity, while model-based rewards, such as ESM-2 sequence perplexity [15], encourage evolutionary plausibility. Unlike DPO’s pairwise preference alignment, PocketX treats these metrics as continuous reward signals, providing richer feedback and finer control in an online RL setting.

Our key contributions are summarized as follows:

- We propose PocketX, a diffusion-based generative model for pocket design that incorporates GRPO to steer generation with biophysical constraints. To the best of our knowledge, this is the first application of GRPO to the diffusion-based generative models for protein design.
- Experiments show that PocketX achieves state-of-the-art performance in generating pockets with more desired properties, such as binding affinity and evolutionary plausibility.

## 2 Methods

In this section, we introduce PocketX, consisting of two training phases, as shown in Figure 1. First, we train a ligand-conditioned generative model [36] to co-design pocket structures and sequences, referred to as the base model (see the section A in the appendix for more details). Next, we fine-tune the base model using the RL algorithm GRPO, where the optimized network learns a policy that simultaneously accounts for biophysical energy and evolutionary plausibility preference. Section 2.1 outlines the preliminaries and notations, and Section 2.2 details the GRPO framework for protein pocket generation.

**Figure 1.**
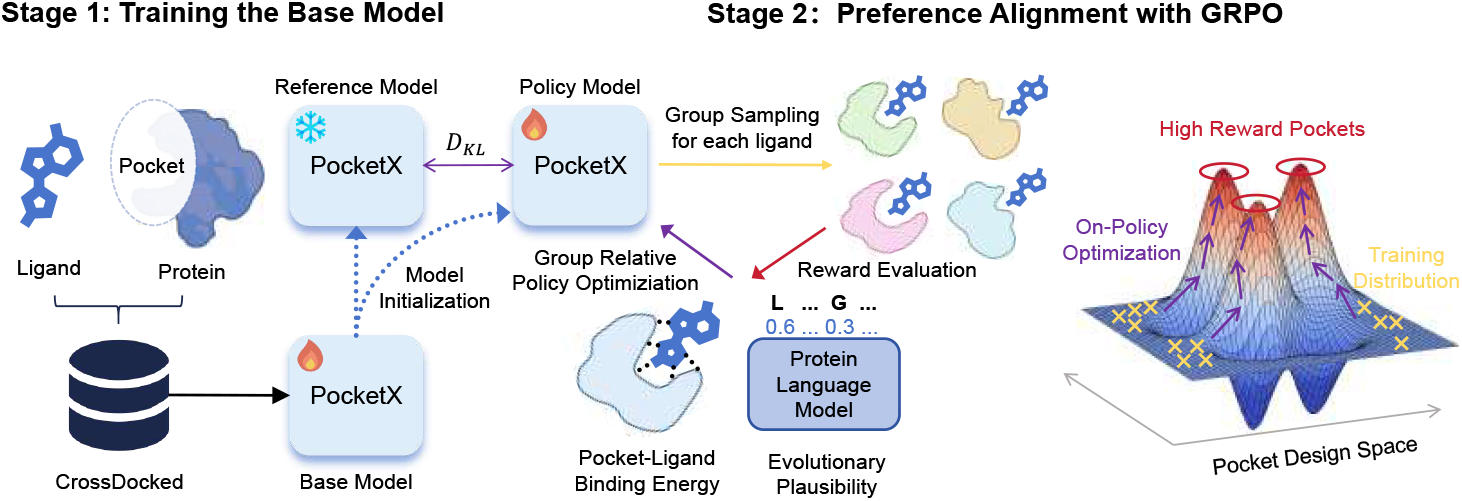
Overview of PocketX. A two-stage framework where a ligand-conditioned base model is trained for pocket structure–sequence co-design, then finetuned with GRPO using biophysical and evolutionary rewards. A reference model regularizes training to avoid over-optimization.

**Figure 2.**
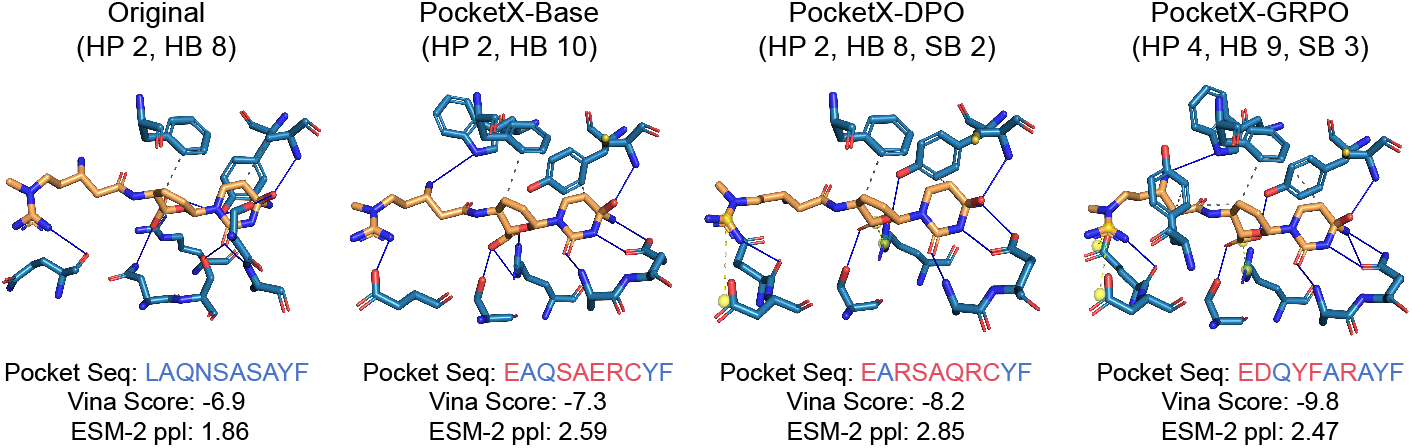
Visualization of protein–ligand interaction analysis for small molecule (PDB ID: 1WN6). HP indicates hydrophobic interactions, HB signifies hydrogen bonds, and SB denotes salt bridges, which are depicted by gray dashed lines, blue solid lines, and yellow dashed lines, respectively. Pocket sequence is shown, with red residues indicating sites that diverge from the native sequence.

### 2.1 Problem Formulation

Following the convention in previous works [46, 47], pocket generation in PocketX is formulated as a conditional generation problem that generates the sequence and structure of the pocket conditioned on the target ligand and the protein scaffold (the protein regions outside the pocket). We model protein–ligand complex as 𝒞 = {𝒫, ℳ, 𝒮}, consisting of the protein pocket 𝒫, protein scaffold 𝒮 and the ligand molecule ℳ. The ligand molecule can be represented as a 3D point cloud 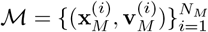, where **x**_*M*_ ∈ ℝ^3^ and **v**_*M*_ ∈ ℝ^*K*^ denote the atomic 3D coordinates and atom type respectively, and *K* being the size of the atom type vocabulary and *N*_*M*_ the number of atoms in the ligand molecule. The protein pocket and protein scaffold are represented as a sequence of residues 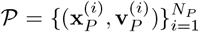 and 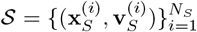, respectively, where *N*_*P*_ and *N*_*S*_ denote the number of amino acids in the protein pocket 𝒫 and the scaffold 𝒮. Note that we represent the 3D atomic positions of a protein residue as **x** ∈ R^14×3^, where 14 is the largest number of atoms of any possible amino acid in the designed protein pocket. With the preceding notations, PocketX is defined as a conditional generative model, formally expressed as *p*_*θ*_(𝒫|𝒮, ℳ).

### 2.2 Group Relative Policy Optimization for Pocket Design

To steer PocketX toward desired biophysical properties, we adopt GRPO [11, 32]. Originally developed for autoregressive language modeling, GRPO’s group-relative updates rely solely on scalar rewards, making it model-agnostic and readily applicable to diffusion and flow models [43, 16]. We apply it to our denoising policy, treating each diffusion trajectory as a rollout and optimizing continuous physics- and model-based rewards online. For each ligand ℳ, the generative model samples a group of *G* individual pockets {𝒫_1_, 𝒫_2_, …, 𝒫_*G*_ from the old policy 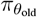 and then optimizes the current policy model *π*_*θ*_ with the training objective:

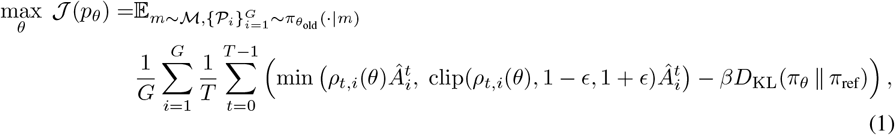

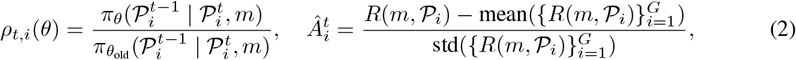

where *t* is the noise level of diffusion process, *ρ*_*t,i*_(*θ*) is the probability ratio, *π*_ref_ is a reference policy, *ϵ* and *β* are hyperparameters, and 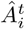 is the advantage, computed using a group of rewards 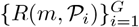 corresponding to the outputs in each group. *D*_KL_(*π*_*θ*_ ∥ *π*_ref_) is the KL divergence between the updated policy and the reference policy. 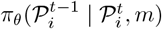 stands for the denoising step, also known as the reverse process:

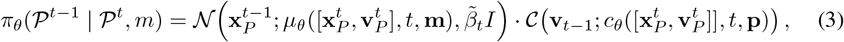

where 𝒩 and 𝒞 denote the Gaussian and categorical distributions, respectively, with their parameters approximated by the model’s predictions *µ*_*θ*_ and *c*_*θ*_.

#### 2.2.1 Reward Design

The whole point of RL is to find a policy that maximizes the expected cumulative reward. Therefore, the reward serves as the training signal of RL, determining the optimization direction [11]. Recognizing the superiority of evolutionary plausibility in protein design [12, 50, 28] and binding energy in drug design [10], we integrate them as reward functions in our RL framework, defined as a weighted combination of evolutionary plausibility and binding energy:

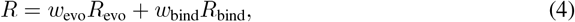

where *R*_evo_ and *R*_bind_ are normalized reward components, and we empirically set their respective weights *w*_evo_ = 0.5 and *w*_bind_ = 1.0.

##### Evolutionary plausibility reward

Evolutionary Plausibility measures how likely a designed sequence is evolutionarily plausible in nature, reflecting adherence to general evolutionary rules of natural [12]. It is evaluated using the likelihood under an independent protein language model. Specifically, we evaluate evolutionary plausibility by calculating the perplexity of the protein language model for the designed pocket region. Here, we use 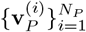 to represent the residue types of the designed pocket 𝒫. The specific formula is formally described as follows:

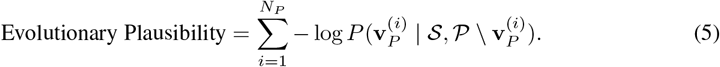

##### Binding energy reward

The binding affinity between the ligand and the receptor protein was evaluated using AutoDock Vina. The Vina scoring function is an optimized, empirical function that provides an estimate of the binding free energy for the ligand-receptor complex. It decomposes the intricate network of intermolecular interactions into a weighted sum of computationally tractable terms. The general form of the scoring function is expressed as:

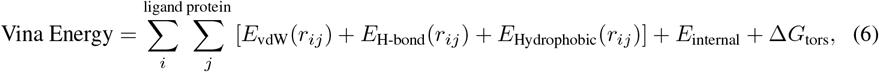

where *i* and *j* index the atoms of the ligand and the protein, respectively. *r*_*ij*_ is the distance between atoms *i* and *j. E*_vdW_(*r*_*ij*_), *E*_H-bond_(*r*_*ij*_), *E*_Hydrophobic_(*r*_*ij*_), and *E*_internal_ represent the van der Waals interaction energy, hydrogen bonding energy, hydrophobic free energy, and the internal energy of the ligand, respectively. Δ*G*_tors_ = *w*_tors_ ·*N*_tors_ is a torsion penalty proportional to the number of rotatable bonds in the ligand (*N*_tors_) restricted upon binding.

## 3 Results

### Datasets

We train and evaluate PocketX on CrossDocked2020 [9], which comprises 22.5M cross-docked protein–molecule pairs. Following prior work [46, 47], we remove samples with binding pose RMSD > 1 Å, yielding ~ 180k data points. For splitting, sequences are clustered at 30% identity using MMseqs2 [34], from which we select 75k pairs for pre-training, 15k pairs for reinforcement learning fine-tuning, and 100 pairs from the remaining clusters for validation and testing. Consistent with previous work [46, 47], the pocket region is defined as the set of all protein residues containing atoms within 3.5 Åof any ligand atom, following the conventional distance ranges pertaining to protein-ligand interactions [19]. Evaluation is performed based on 100 independently sampled pockets for each ligand in the test set. More details about implementation can be seen in section B.

### Evaluation Metrics

We adopt the same evaluation metrics as prior work [46, 47] to assess both the sequence and structural validity of generated pockets. Amino Acid Recovery (**AAR**) measures the sequence recovery accuracy by comparing the generated sequences to the native sequences. Self-consistency Root Mean Squared Deviation (**scRMSD**) measures the deviation between the generated backbone atoms and the predicted pocket’s backbone, serving as an indicator of structural plausibility. Specifically, for each generated protein structure, eight sequences are derived by ProteinMPNN [4] and then folded to structures with ESMFold [15]. Binding energy is evaluated with **AutoDock Vina** [8] and **GlideSP**, whereas evolutionary plausibility is measured by the likelihood under an independent protein language model **ESM-2** [15].

### Baseline Methods

We evaluate PocketX in comparison with representative methods from each category: the traditional method PocketOptimizer [20], and three recent deep learning-based models FAIR [46], RFdiffusion All-Atom [14], and PocketGen [47]. More details on running these methods are in Appendix C

### Experimental Results

As shown in Table 1, Fig. 3, and Fig. 4, PocketX-GRPO achieves the lowest binding energies while maintaining competitive structural validity and evolutionary plausibility. These consistent improvements over PocketX-Base and PocketX-DPO demonstrate that GRPO is an effective optimization paradigm, steering generation under biophysical constraints.

**Table 1:** Evaluation on CrossDocked2020 test set. Here, *reference* represents the native pocket structure and sequence in the dataset. *SR* denotes the sidechain relaxation for the additional post-processing. We highlight the best two results with **bold text** and <u>underlined</u> text, respectively. The scRMSD results for PocketOpt are omitted, as the method keeps the protein backbone structures fixed. ∗ indicates that *SR* only affects sidechain conformations.

| Methods | Binding Energy<br>Vina Score (↓) | Binding Energy<br>GlideSP (↓) | Structure Validity<br>scRMSD (↓) | Sequence Plausibility<br>AAR (↑) | Evolutionary Plausibility<br>ESM-2 ppl (↓) |
| --- | --- | --- | --- | --- | --- |
| *reference* | -7.17 | -6.60 | 0.65 | - | 2.36 |
| PocketOpt | -6.86 | -6.33 | - | 25.86% | 2.95 |
| FAIR | -7.01 | -6.45 | 0.85 | 14.55% | 3.34 |
| RFdiffusionAA | -6.93 | -6.42 | 1.16 | **37.62%** | **2.49** |
| PocketGen | -7.19 | -6.64 | 0.79 | 14.83% | 3.31 |
| PocketGen *w/o SR* | -6.48 | -6.19 | * | * | * |
| PocketX-Base | -7.21 | -6.69 | **0.67** | 28.9% | 2.87 |
| **PocketX-DPO** | <u>-7.87</u> | <u>-7.16</u> | 0.80 | 27.21% | 2.93 |
| **PocketX-GRPO** | **-8.43** | **-7.85** | <u>0.68</u> | <u>34.89%</u> | <u>2.56</u> |

**Figure 3.**
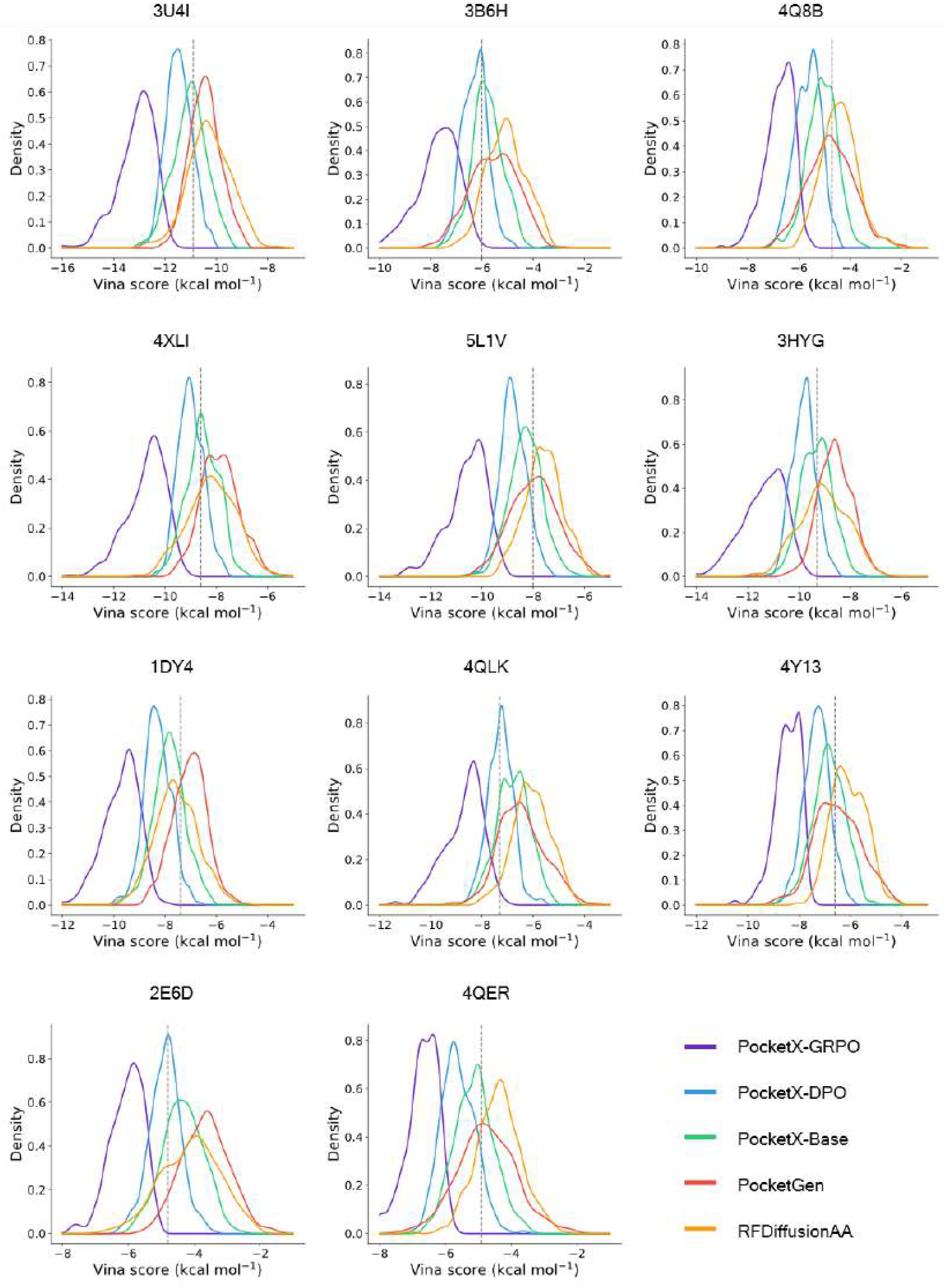
Pocket binding affinity distributions of PocketX and baseline methods for the target molecules in PDB. We mark the Vina Score of the original pocket with the vertical dotted lines. For each method, we sample 100 pockets for each target ligand.

**Figure 4.**
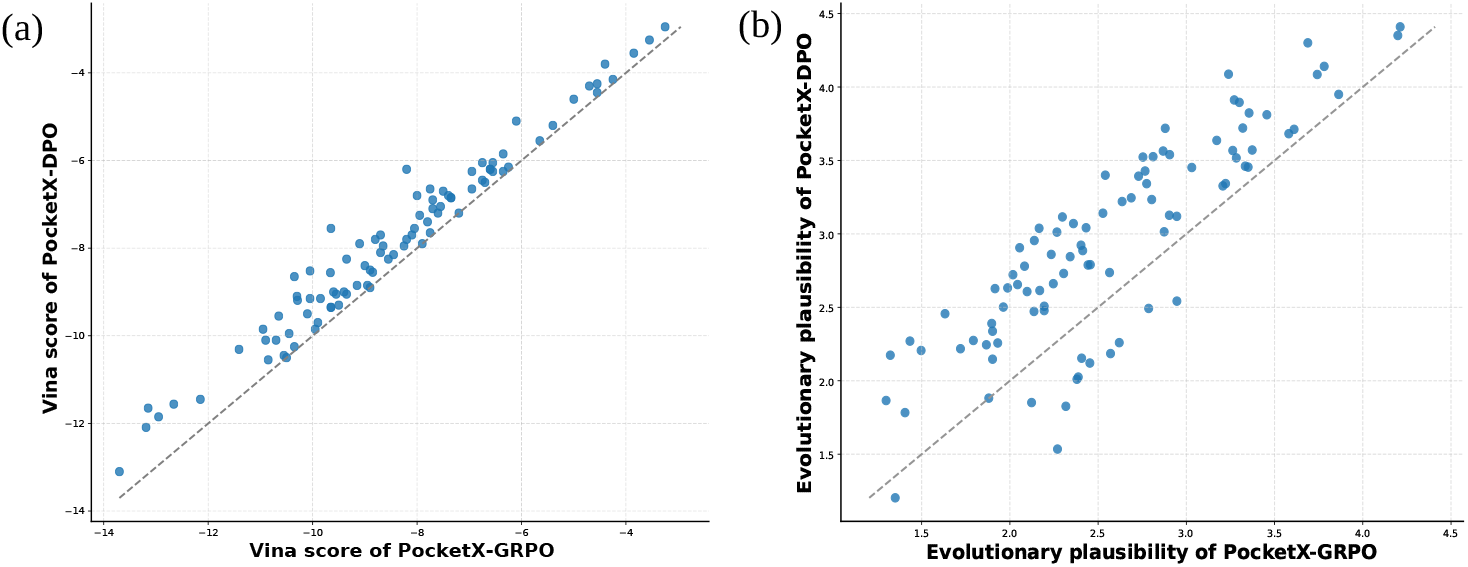
Head-to-head comparison across reward signals. Left: AutoDock Vina score; Right: Evolutionary plausibility.

### Interaction Analysis

As shown in Fig. 2, we use PLIP [30] to analyze the pocket–ligand interactions and compare them with the native pattern. Across different variants, PocketX introduces additional physically plausible contacts; notably, GRPO fine-tuning (vs. DPO) enriches hydrophobic and electrostatic interactions—e.g., via Phenylalanine and Aspartic acid—leading to more favorable binding energy and evolutionary plausibility, demonstrating the advantage of GRPO.

## 4 Conclusions

We propose PocketX, a diffusion-based generative model that integrates online reinforcement learning for protein pocket design. Using Group Relative Policy Optimization (GRPO), PocketX consistently steers generation toward pockets that satisfy the desired biophysical properties, as confirmed by our experiments. The framework is modular and readily accommodates additional reward signals. For example, predicted confidence scores from AlphaFold3 [1] and affinity estimates from Boltz-2 [22] could be incorporated in an online RL setting as the guidance of protein-ligand complex structures and binding affinity. Exploring these extensions is a central focus of our future work and is expected to enhance PocketX’s performance even further.

## Acknowledgments and Disclosure of Funding

We acknowledge the financial support from the National Key R&D program of China (grant no. 2023YFF1205103) and the National Natural Science Foundation of China (grant no. 32370657).

## A Details of Diffusion Processes

For the completeness of our study, we provide a brief introduction to diffusion and model architectures of the denoiser.

### A.1 Diffusion Process for Sequence

For the protein sequence, we employ *absorbing* discrete diffusion framework [2, 48], a widely used diffusion model in protein design [40, 41]. Let Cat(**v**) be a categorical distribution on sequence **v**. The forward process of discrete diffusion defines a Markov process governed by the transition kernel

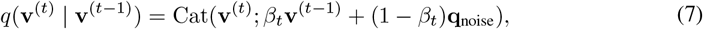

where 0 *β*_*t*_ ≪ 1 is the noise schedule controlling the degree of corruption at timestep *t*, and it gradually perturbs the data **v**^(0)^ ~ *q*(**v**^(0)^) into a stationary distribution **v**^(T)^ ~ **q**_noise_. For absorbing diffusion, **q**_noise_ is the point mass with all of the probability on the mask state.

### A.2 Diffusion Process for Structure

For the protein structure, we employ the EDM [13] framework to model the diffusion process. Let us denote the protein structure distribution by *q*(**x**^(0)^), with standard deviation *σ*_struc_data_, and consider the family of mollified distributions *q*(**x**^(*t*)^; *σ*_*t*_) obtained by the forward process

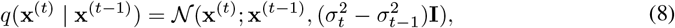

where 𝒩 is Gaussian distribution. For *σ*_*t*_ ≫ *σ*_struc_data_, *q*(**x**^(*t*)^; *σ*_*t*_) is practically indistinguishable from pure Gaussian noise.

### A.3 Details of Model Architectures

While previous protein generators typically use small equivariant neural networks, we take inspiration from AlphaFold3 [1] and utilize a non-equivariant network. Our model comprises two primary components: the InputEmbedder module and the Diffusion module. The Diffusion module consists of an AtomTransformer encoder, a Diffusion transformer, and an AtomTransformer decoder. While all three components are constructed as transformer stacks, the AtomTransformer encoder and decoder operate on atom-level representations, whereas the DiT [23] operates on token-level representations. At each time step t, the network receives the noisy sequence **v**^(*t*)^ and structure **x**^(*t*)^ as inputs, and predicts the denoised sequence **v**^(0)^ and structure **x**^(0)^.

## B Details of Implementation

### B.1 Pretraining Details

For PocketX-Base model training, we adopted the DeepSpeed [27] strategy and used the Adam optimizer [17] with a learning rate of 0.0001 and parameters *β* values of (0.9, 0.999). The pretraining was conducted on 4 NVIDIA 80G H100 GPUs with a batch size of 64 and a gradient norm clipping value of 1.0, and achieved convergence within 100k steps.

### B.2 Reinforcement Learning Details

For DPO and GRPO reinforcement learning, the pretrained base model was optimized using the Adam optimizer with an initial learning rate of 5 × 10^−5^ and *β* = (0.95, 0.999). All other hyperparameter settings were kept consistent with those used in pretraining. Following previous works [10, 49], we construct preference-optimized datasets for pocket and ligand complexes. We generated 128 pockets for each ligand in the fine-tuning dataset, and the reward metrics, such as binding energy and evolutionary plausibility, are calculated for these pockets. The highest-scoring samples under the reward function were treated as preferred, in contrast to the lowest-scoring samples, which were treated as dispreferred. Training of the PocketX-DPO model was conducted on 4 NVIDIA H100 GPUs, achieving convergence in 30k steps. We configured the group size as *G* = 8 for the reinforcement learning process of GRPO, generating eight candidate protein pockets for each ligand molecule. This setting offered a reliable optimization signal while maintaining acceptable computational overhead. The KL ratio *β* in the GRPO loss function is set as 0.05. The PocketX-GRPO model was trained on 4 NVIDIA H100 GPUs and converged within 80k steps.

## C Details of Baseline Methods

**PocketOptimizer [20]** is a physics-based computational approach for protein design, which predicts mutations within protein binding pockets to enhance affinity for a target ligand. In this study, we employ its most recent release, PocketOptimizer 2.0. The typical workflow of PocketOptimizer involves four key stages: preparing the protein structure, sampling conformational flexibility, calculating energetic contributions, and generating candidate design solutions. During the energy evaluation step, both packing energies and ligand-binding energies are taken into account. In line with the original protocol, the protein backbone was held fixed throughout the design process. We use the test script provided in the GitHub repository (https://github.com/Hoecker-Lab/pocketoptimizer). All the hyperparameters we used are default.

**RFdiffusion All-Atom [14]** is the latest iteration of RFDiffusion, combining residue-level representations for amino acids with atomic representations for other molecular groups. It supports modeling diverse molecular complexes, such as proteins with small molecules, metals, nucleic acids, or covalent modifications. Starting from the random noise of residues around target molecules, it can directly generate the binding protein backbone. However, to complete protein design, especially pocket design, explicit residue identities are required. Thus, LigandMPNN [6], the recently updated version of ProteinMPNN [4], is employed to predict the amino acid type at each position. We use the pretrained model and test script provided in the GitHub repository (https://github.com/baker-laboratory/rf_diffusion_all_atom). All the hyperparameters are default.

**FAIR [46]** is an algorithm for the co-design of pocket sequences and structures. It performs a two-stage process in a coarse-to-fine fashion, starting with backbone refinement and proceeding to full-atom refinement, including side chains, to generate full-atom pockets. We employed FAIR from its GitHub repository (https://github.com/zaixizhang/FAIR) with all default hyperparameters.

**PocketGen [47]** is a deep generative method developed for efficient protein pocket design. It employs a co-design approach in which both the sequence and structure of a protein pocket are predicted from the ligand and the surrounding protein scaffold (excluding the pocket itself). The model architecture consists of two main components: a bilevel graph transformer and a sequence refinement module. We use the pretrained model checkpoint provided in the GitHub repository (https://github.com/zaixizhang/PocketGen). All the hyperparameters we used are default.

